# Toward overcoming pyrethroid resistance in mosquito control: the role of sodium channel blocker insecticides

**DOI:** 10.1101/2023.03.29.534712

**Authors:** Beata Niklas, Jakub Rydzewski, Bruno Lapied, Wieslaw Nowak

## Abstract

Diseases spread by mosquitoes lead to death of 700,000 people each year. The main way to reduce transmission is vector control by biting prevention with chemicals. However, the most commonly used insecticides lose efficacy due to the growing resistance. Voltage-gated sodium channels (VGSCs), membrane proteins responsible for the depolarizing phase of an action potential, are targeted by a broad range of neurotoxins, including pyrethroids and sodium channel blocker insecticides (SCBIs). Reduced sensitivity of the target protein due to the point mutations threatened malaria control with pyrethroids. Although SCBIs – indoxacarb (a pre-insecticide bioactivated to DCJW in insects) and metaflumizone – are used in agriculture only, they emerge as promising candidates in mosquito control. Therefore, a thorough understanding of molecular mechanisms of SCBIs action is urgently needed to break the resistance and stop disease transmission. In this study, by performing an extensive combination of equilibrium and enhanced sampling molecular dynamics simulations (3.2 μs in total), we found the DIII-DIV fenestration to be the most probable entry route of DCJW to the central cavity of mosquito VGSC. Our study revealed that F1852 is crucial in limiting SCBI access to their binding site. Result explain the role of the F1852T mutation found in resistant insects and the increased toxicity of DCJW compared to its bulkier parent compound, indoxacarb. We also delineated residues that contribute to both SCBIs and non-ester pyrethroid etofenprox binding and thus could be involved in the target site cross-resistance.

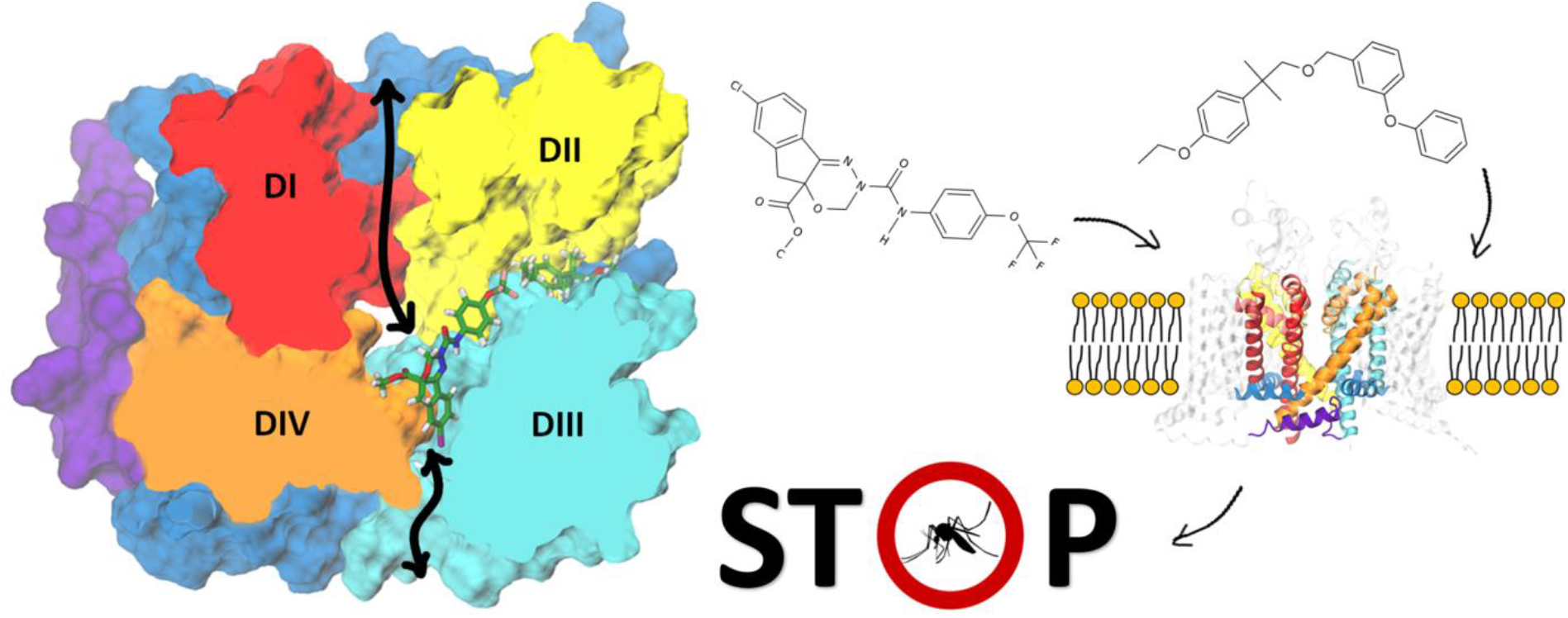

## 1. INTRODUCTION

The World Health Organization (WHO) warns that more than half of the human population is currently at risk of mosquito-borne diseases. It is predicted that progressive climate change will increase the extent of outbreaks [1, 2]. Malaria is a life-threatening disease caused by *Plasmodium* parasites transmitted to people through the bites of infected female *Anopheles* mosquitoes. In 2021, an estimated 247 million malaria cases led to approximately 619 000 deaths, with a tremendous toll on children under 5 years old (WHO, World Malaria Report 2022).

The primary way to reduce transmission is biting prevention with chemicals. Vector control relies heavily on insecticides and thus can be compromised by resistance [3, 4]. Resistance to pyrethroids, the only insecticide class integrated into bed nets, is widespread globally [5]. To find new ways to control insect populations, a thorough understanding of molecular mechanisms of insecticide action and resistance is required.

The voltage-gated sodium channels (VGSCs) represent one of the major molecular targets of insecticide action. These multi-domain transmembrane proteins are responsible for the depolarizing phase of action potentials in nerves and muscles [6]. While nine subtypes of VGSCs are expressed in humans (hNav1.1-hNav1.9), a single copy of gene coding a sodium channel protein (~2,050 amino acids) can be found in most insects [7, 8]. The α-subunit comprises four homologous domains (DI-DIV) including six transmembrane helices (S1-S6) each (Figure 1a). Helices S1-S4 constitute the voltage-sensing domain (VSD) with a positively charged S4 helix acting as a voltage sensor, while helices S5 and S6 linked by membrane-reentrant pore loop (P-loop) contribute to the ion-conducting pore domain (PD). Due to the crucial role of VGSCs in regulating neuronal membrane excitability, they are targeted by a broad range of naturally occurring and synthetic neurotoxins that alter sodium conductance by blocking the ion-conducting pore or altering gating (opening and closing of a channel) [9]. Some of them – DDT, pyrethroids, and sodium channel blocker insecticides (SCBIs) – are known for their insecticidal activity [10].

**Figure 1.**
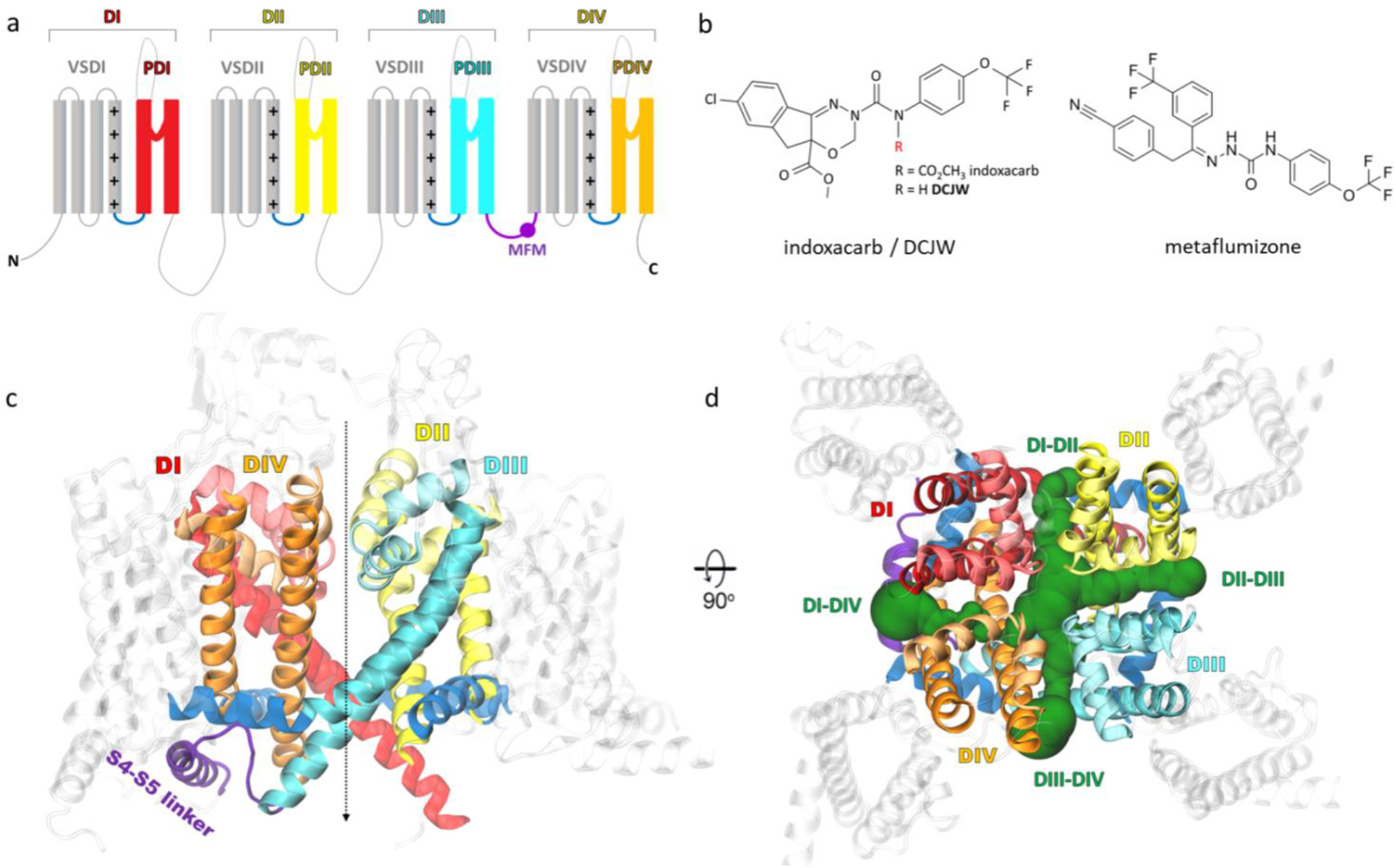
a) Topology of the pseudotetrameric α-subunit of the mosquito *Anopheles gambiae* voltage-gated sodium channel shows four domains consisting of six transmembrane helices each (S1-S6). Helices 1-4 contribute to the voltage-sensor domain (VSD, grey), with a positively charged S4 helix acting as a voltage sensor, and helices S5-S6 create the pore domain (PD). The S4-S5 linkers are shown in dark blue. The inactivation gate MFM motif, as a part of the DIII-DIV linker, is marked in violet. b) Structures of sodium channel blocker insecticides: indoxacarb/DCJW (left) and metaflumizone (right). c) Homology model of the mosquito channel based on the inactivated-state structure of the hNav1.7 channel (PDB code: 6J8G [11]). The approximate location of the pore is indicated by the dashed line. d) The side (left) and top (right) view of the channel with the four lateral fenestrations found by MOLE shown in green.

The extensive use of pyrethroids for the last 50 years has led to the development of knockdown resistance (kdr) in insects worldwide. Over 50 mutations that reduce neuronal sensitivity to DDT and pyrethroids were reported in the α-subunit of VGSC in arthropod species [8]. Due to the reduced sensitivity of target protein by point mutation and/or metabolic resistance (i.e., overexpression of detoxifying enzymes), the effectiveness of long-lasting insecticidal nets and indoor residual spraying treatment of houses with pyrethroid formulations are under threat. No alternative is recommended by WHO Pesticide Evaluation Scheme (WHOPES) for use on mosquito nets. Therefore, there is an urgent need to develop new chemicals for efficient vector control.

SCBIs are relatively new and highly selective insecticides. Two compounds of this group – indoxacarb and metaflumizone - were commercialized [12]. These potent inhibitors of VGSCs have an overlapping binding site with local anesthetics (e.g., lidocaine) [12]. SCBIs preferably bind to and trap VGSC in the slow-inactivated, nonconducting state and block neuronal action potentials in both peripheral and central nervous systems [13, 14].

Indoxacarb (Figure 1b), an oxadiazine SCBI, was developed in 1992 by DuPont by optimizing pyrazoline analogs to limit non-target activity in mammals and other organisms and to increase environmental safety while maintaining insecticidal efficacy [15]. It was registered in 2000 as a reduced-risk compound. In insects, indoxacarb undergoes rapid bioactivation to the more neurotoxic DCJW (N-decarbomethoxyllated JW062) derivative [14], leading to uncoordinated movement, tremors, and pseudoparalysis - a state characterized by violent convulsions in paralyzed insects when disturbed. The differences between mammalian and arthropodic metabolism of indoxacarb partially account for its selectivity towards insects and the consequent widespread usage in agriculture.

Metaflumizone (Figure 1b), a semicarbazone SCBI, was registered to use on Chinese cabbage fields in 2009 as it provides high-efficiency control of the most economically important pests. Formulated with amitraz, metaflumizone controls fleas, ticks, and mites on dogs and cats [16]. It is considered a low-risk chemical to non-target organisms, including pollinators, safe for mammals and the environment [17].

To this date, all the active ingredients used for malaria vector control are spin-offs from agricultural uses [18]. Screening existing registered pesticides provides a rapid route to identifying chemicals of potential value to public health. In the evaluation of 81 commercial agrochemicals for mosquito control, metaflumizone, acetamiprid, thiamethoxan, and thiocyclam were the most promising candidates [18]. Both metaflumizone and indoxacarb were selected in a testing cascade against adult female *Anopheles gambiae* mosquitoes as compounds of high interest as vector control agents [19]. In tunnel tests, indoxacarb induced delayed mortality over 24-96 h against both pyrethroids-sensitive and resistant *An. gambiae,* but there was no protection regarding blood-feeding inhibition [20]. Neither synergism nor antagonism was observed between indoxacarb and pyrethroid [20]. In further tunnel tests and bioassays, indoxacarb was highly effective in both mortality and reducing the blood feeding by a pyrethroid-susceptible strain of *An. gambiae* and pyrethroid-susceptible and resistant strains of *C. quinquefasciatus* mosquitoes [21]. Mixtures of indoxacarb with pyrethroid alphacypermethrin conferred far greater protection than the single treatments for mortality and blood-feeding inhibition [21]. Thus, the combination of SCBIs and pyrethroids is certainly worth further investigation in terms of both efficacy in vector control and potential cross-resistance. Resistance may involve multiple mechanisms within the insect, e.g., cuticular permeability, metabolic degradation, behavioral resistance or point mutation in the target protein [22]. Although SCBIs and pyrethroids bind to distinct sites on VGSC, mutagenesis data suggest that their binding regions may partially overlap [23]. It is necessary to investigate the role of the target-site in the development of cross-resistance between pyrethroids and SCBIs.

The hydrophobic access pathway for small molecules from the plasma membrane to the center of the ion-conducting pore referred to as the central cavity (CC) of VGSC is *via* four lateral fenestrations (Figure 1d) [24]. These tunnels are delineated by interfaces between the S5 and S6 helices of two adjacent PDs, named DI-DII, DII-DIII, DIII-DIV, and DI-DIV. Multiple structures of eukaryotic VGSC have shown that fenestrations are not symmetrical. Their size changes during the gating cycle and differs across VGSC subtypes [25]. A hypothetical pathway of metaflumizone to the central cavity of VGSC *via* DIII-DIV fenestration was proposed [26]. However, the dynamical modeling of SCBIs’ entrance to the central cavity of VGSC was not provided yet. DCJW has a rigid, tricyclic core (Figure 1b), which makes the ligand entrance to the CC of the channel through the fenestration questionable. It is necessary to estimate to what extent fluctuations of the fenestration shape enable the access of SCBIs to modulate VGSC and thus insect physiology.

Ligand-protein association is known to occur on timescales far exceeding the capabilities of equilibrium molecular dynamics (MD) simulations [27, 28]. Therefore, in this study, we perform an extensive combination of equilibrium and enhanced MD simulations of a total aggregate simulation time of 3.2 μs to alleviate the sampling problem [29] and to find the entry route of DCJW to the CC of mosquito VGSC. We investigate the binding interactions of DCJW, metaflumizone, and a non-ester pyrethroid etofenprox within the AgNav1 channel to assess the impact of target site insensitivity conferring resistance to pyrethroids (kdr) on SCBIs action on mosquito channel.

## 2. RESULTS AND DISCUSSION

### 2.1 Docking of DCJW and Metaflumizone to the *An. gambiae* VGSC

The binding of SCBIs to the slow-inactivated channel state is favored over the open state not due to the conformation of the VGSC itself but rather due to the very slow kinetics of the channel-ligand association [30]. This is supported by the observation that upon removing slow and fast-inactivation, the block of fast-inactivated and open channels by the pyrazoline RH-1211 proceeds as quickly as the block of slow-inactivated channels [31]. Bearing this in mind, we built three models of mosquito AgNav1 VGSC: open, inactivated, and closed, based on the templates of experimental structures captured in those states [11, 32, 33].

We found that in the lowest energy docking poses in both open- and inactivated-state models, DCJW and metaflumizone extend between the DIII-DIV fenestration, CC, and the entrance to the DII-DIII fenestration from the interior of the channel. The root-mean-square deviation (RMSD) values between the open- and inactivated-state models are 1.7 Å for DCJW and 2.81 Å for metaflumizone. The binding energy of docking to the inactivated model, measured by the smina scoring function (SSF), equals −9.61 kcal/mol for DCJW and −10.18 kcal/mol for metaflumizone (Figure 2). We found no poses of either DCJW or metaflumizone in the PD of the closed AgNav1 model. However, in their lowest energy poses (SSF=-8.05 kcal/mol for DCJW and −8.42 kcal/mol for metaflumizone), both ligands approach the DIII-DIV fenestration from the plasma membrane; see Figure S1 in the Supporting Information (SI). As the inactivated-state model (Figure 1c,d) reflects the most physiological binding condition, we focus on this model further.

**Figure 2.**
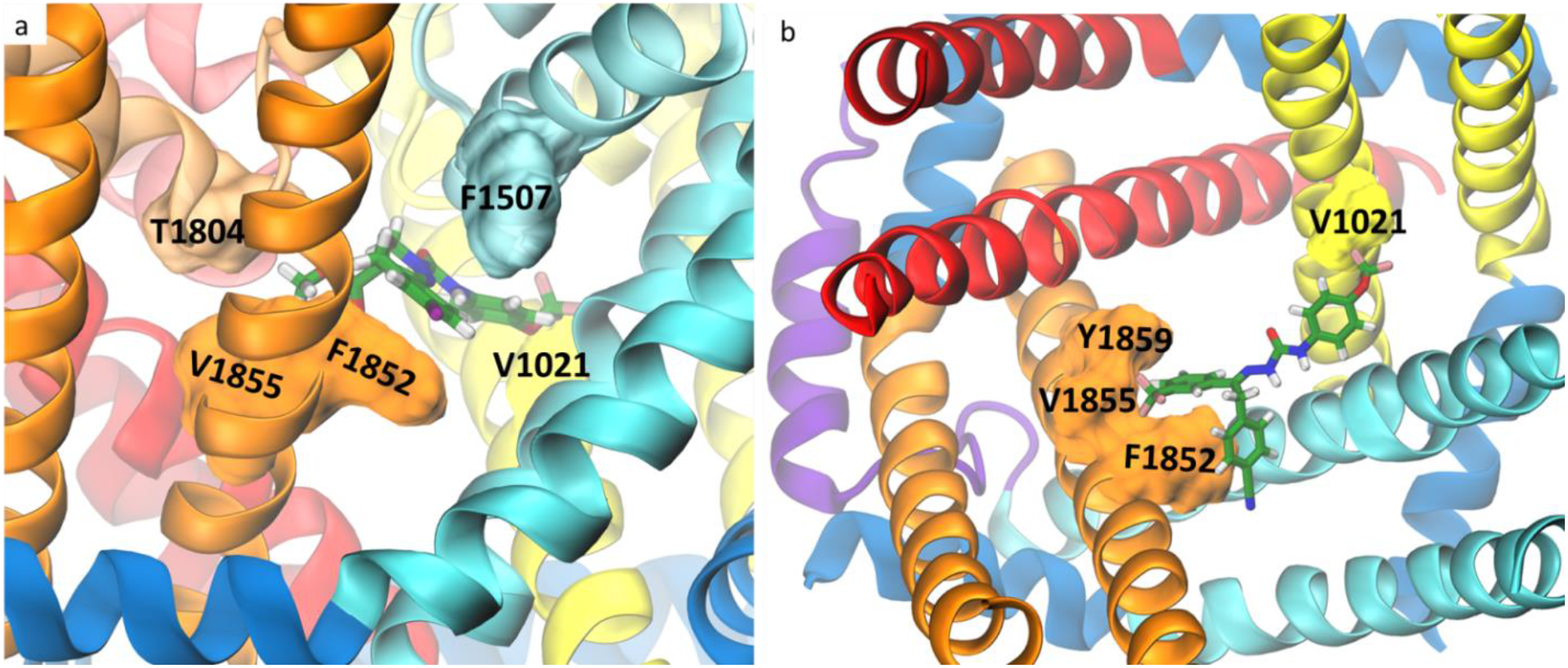
Docking of DCJW (a) and metaflumizone (b) to the mosquito voltage-gated sodium channel (VGSC). Residues known to affect channel sensitivity to these insecticides are shown in a surface representation. Ligands are shown in a stick representation with carbon atoms in green, nitrogen in blue, oxygen in red, fluor in pink, chloride in purple, and hydrogen atoms in white. Side view is presented in (a) and top view with the P-loop removed for clarity in (b). Coloring of the channel helices refers to Figure 1.

A common structural backbone for SCBIs, which may play a key role in their insecticidal activity, consists of two aromatic rings connected by five atoms including three nitrogens and one carbonyl group [34]. In our docking, nitrogen atoms from both ligands form hydrogen bonds with S1552 from the DIII S6 helix, while the carbonyl group faces the channel axis to potentially block the flow of the sodium ions through the pore. The highest contribution to the docking binding energy of both DCJW (21%) and metaflumizone (22%) comes from π-stacking interactions with F1852, which has been found in SCBIs-resistant insects [35, 36]. Further contacts are primarily hydrophobic, and most of them include residues from S6 helices and P-loops from DIII and DIV. Among them, we found F1507, T1555, T1804, V1855, and Y1859, the substitution of which affected SCBIs activity on VGSCs [26, 37, 38]. While the trifluoromethyl group of DCJW approaches but does not penetrate the DII-DIII fenestration, the trifluoromethyl group of metaflumizone is buried in the tunnel. Both ligands interact with V1021 from DII S6, the substitution of which was found to affect the rNav1.4 channel sensitivity to indoxacarb and metaflumizone but not DCJW [39]. To validate this result upon fluctuating channel conformations, we run three 250 ns MD simulations of DCJW-bound AgNav1. In all three MD simulations, DCJW stayed tightly bound and its energy (rescored after the simulations using the docking function SSF) reached −10.96 kcal/mol due to more favorable side chain orientations. We found a hydrogen bond between S1552 and the ligand in >50% of all snapshots. We also observed that the sodium ion and the carbonyl group of DCJW stayed close to each other in all three MD simulations (Figure S2 in the SI). For a list of the channel-DCJW contacts observed in our MD simulations, see the SI Table S1.

### 2.2 DCJW Binding Pathway to the Central Cavity

As protein tunnels can be transient and heterogeneous, which may lead to spontaneous closing and opening [40, 41], we aimed to find possible entry and exit routes for DCJW using MD simulations instead of analyzing only the static structures of VGSC. To assess fluctuations of the fenestrations and compare their bottleneck radius distributions, we ran three 250 ns unbiased MD simulations of the inactivated-state AgNav1 model. From Figure 3a, we can see that the DI-II and DIII-DIV fenestrations are the widest, with average bottleneck radii of around 2 Å, making them the most accessible and probable entry route for ligands. This result agrees with a similar analysis performed on human Nav1.1, Nav1.2, Nav1.4, Nav1.5, and Nav1.7 inactivated-state VGSCs [25]. Then, we ran 30 enhanced sampling MD simulations in which we biased the positions of DCJW to search for possible channel entry and exit routes. We found that DCJW could migrate outside the channel only through the DI-DII, DII-DIII, and DIII-DIV fenestrations. For the DI-DIV tunnel, we did not observe any dissociation event (i.e., pathways). As the DII-DIII fenestration is less likely to serve as the entry route for DCJW to reach the CC (Figure 3a), we focused our further investigation on the DI-DII and DIII-DIV fenestrations.

**Figure 3.**
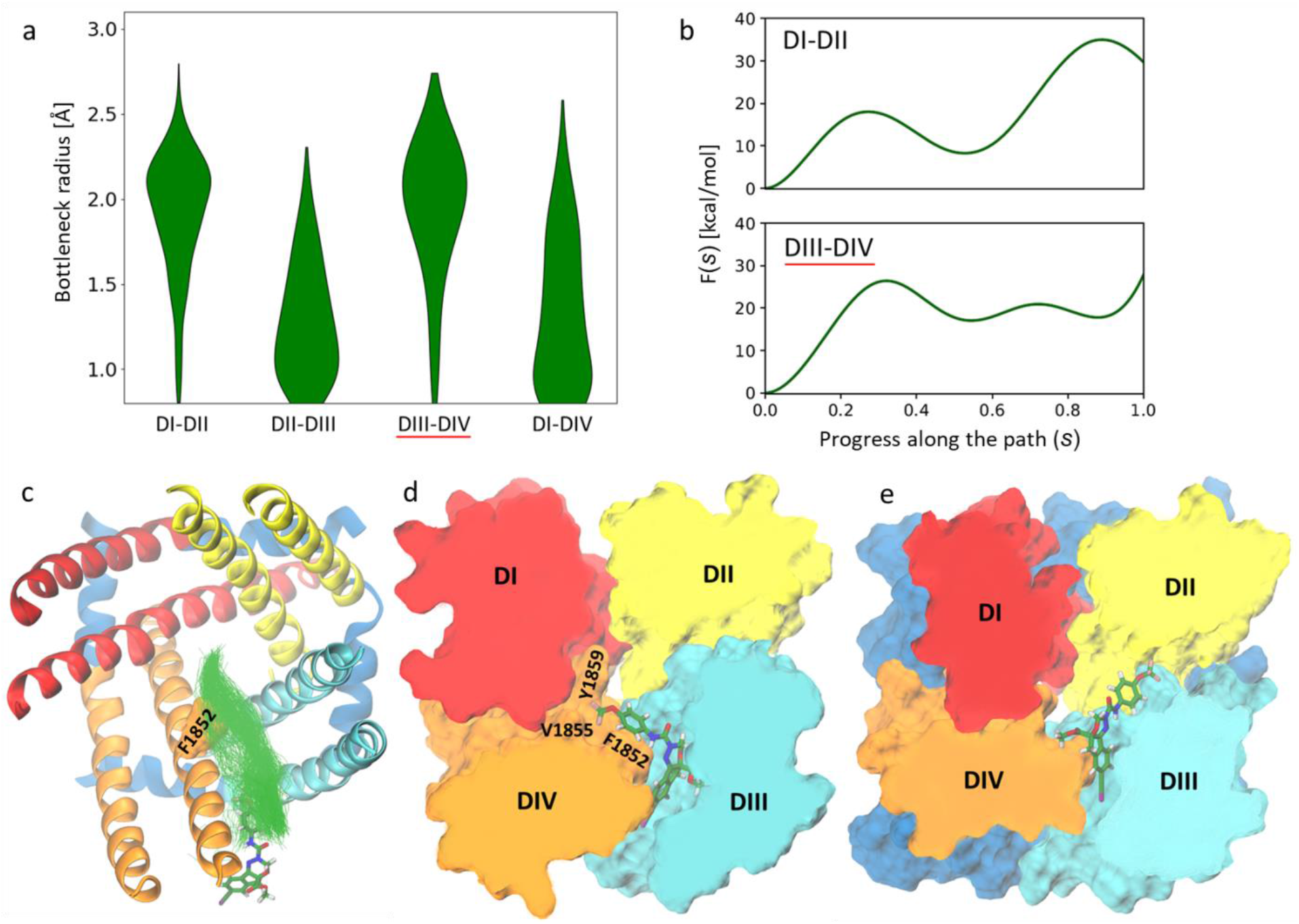
a) The bottleneck radius distributions of the four lateral fenestrations in the inactivated-state model of the mosquito sodium channel calculated using CAVER3.0 [43] based on three 250 ns MD trajectories combined. b) Free energy profiles of DCJW crossing the DI-DII (top) and DIII-DIV (bottom) fenestration. The path starts (0.0) at the DCJW binding site found in docking (shown in e) and ends (1.0) at the channel-membrane interface (indicated by a ligand in sticks representation in c). c) Tunnel clusters found by CAVER in the metadynamics simulations of DCJW crossing the DIII-DIV fenestration are presented as green lines. d) The representative snapshot of the most densely populated cluster of ligand positions along the binding pathway, corresponding to the local energy minimum, where DCJW interacts with Y1852, V1855, and Y1859, known to affect SCBIs’ efficacy on VGSCs.

As the binding reaction coordinates, we selected DCJW conformations from the shortest trajectories that ended in dissociation along the DI-DII and DIII-DIV fenestrations. Next, we estimated the free-energy profiles by biasing path-collective variables [42], i.e., the progress along (s) and distance from the identified binding reaction coordinates (z) along the DI-DII and DIII-DIV fenestrations, during which DCJW could move between the associated and dissociated states in both directions. For each binding reaction coordinate, we performed a 500 ns well-tempered metadynamics simulation. During these enhanced sampling simulations, we observed many transitions across the path-collective variables, suggesting that the simulations converged.

The free-energy profiles as functions of the progress along the DI-DII and DIII-DIV fenestrations, are shown in Figure 3b. Entering the channel, DCJW encounters a free-energy barrier of about 20 kcal/mol when approaching the DIII-DIV fenestration (Figure 3c) and a barrier higher than 30 kcal/mol in the case of the DI-DII fenestration (Figure 3b). Moreover, while the profile along the DII-DIII tunnel is flat, the ligand can get stuck in the free-energy minimum observed in the middle of the pathway *via* the DI-DII fenestration. Also, as the DCJW binding site is located at the intersection of the CC and the DIII-DIV fenestration, the route *via* this tunnel is shorter when compared to the pathway through DI-DII, which includes traveling through the middle of the ion-conducting pore. These observations suggest that the entry to the CC of VGSC through the DIII-DIV fenestration is preferable. Interestingly, after reaching the CC, the ligand faces a high free-energy barrier of about 30 kcal/mol to unbind, which can be explained by a high affinity of DCJW-channel interactions that stabilize the ligands in its binding site (Figure 3e). The height of free-energy barriers seen along the DCJW pathway in both fenestrations agrees well with the very slow kinetics of pyrazolines entry to the binding site and their inability to block open channels [31].

The dynamical variability in the bottleneck radius of fenestration conditionalizes the entry of pore blockers to the CC with the kinking at the midpoint of S6 helices, identified as important in channel gating [44], appearing to be restrictive [25]. In the AgNav1 channel, F1852 is the key hydrophobic residue bottlenecking the DIII-DIV fenestration (Figure 2 and Figure 3c,d). Substitution of the corresponding residue to the F1852 by a bulkier tyrosine had been found in SCBIs-resistant populations of the tomato leafminer and the diamondback moth [35, 36]. The substantial reduction of VGSC sensitivity to indoxacarb due to F1852Y mutation has also been functionally validated by electrophysiological studies in *Xenopus* oocytes [37]. Furthermore, the alanine substitution (F1817A in the cockroach BgNa_v_) enhanced the ability of both DCJW and metaflumizone to interact with inactivated VGSCs. It also provided an easier escape route for metaflumizone to leave its receptor site during recovery from inactivation [38]. We suggest that the resistance could be explained by the reduced frequency of reaching the binding site due to the higher free-energy barrier of crossing the DIII-DIV fenestration.

Interestingly, in docking of indoxacarb, the proinsecticide that undergoes bioactivation to the more toxic DCJW, we obtained the lowest energy poses in the DIII-DIV fenestration (SI Figure S3). We postulate that the higher probability of passing the fenestration due to a decrease of the ligand size upon metabolization may partially explain the increased toxicity of DCJW when compared to indoxacarb.

### 2.3 Comparison of the SCBIs and etofenprox binding region. Implications for the target site cross-resistance with pyrethroids

Recently, high cross-resistance with deltamethrin, the most important pyrethroid in malaria prevention, was observed in an indoxacarb-resistant population of the fall armyworm, *Spodoptera frugiperda* [45]. Little positive cross-resistance between indoxacarb and pyrethroids was also found in the diamondback moth, *Plutella xylostella* L. [46–48]. Negative cross-resistance observed in fenvalerate- and cypermethrin-selected populations of *Helicoverpa armigera* was explained by the elevated level of carboxyl esterase in pyrethroids-resistant insects, which may increase the enzymatic activation of indoxacarb to more toxic DCJW derivative [49]. It is in agreement with a study on the same species, where high resistance to cypermethrin and deltamethrin but not to the non-ester pyrethroid etofenprox and indoxacarb, was observed together with the positive correlation with esterase activity. The lack of resistance to DDT, which acts on the same site as pyrethroids, excludes the involvement of target site resistance in this population [50]. Contradicting results suggest multiple mechanisms involved in indoxacarb resistance. Clearly, there is a shortage of data describing the involvement of the target-site sensitivity in cross-resistance development.

Etofenprox (Figure 4a) is a pyrethroid derivative with an ether linkage instead of an ester linkage present in traditional pyrethroid insecticides, making it less prone to metabolic degradation by esterases. Experiments performed with heterologous VGSC expression in the *Xenopus* oocyte suggest that etofenprox displays a pyrethroid-like effect [51]. It exhibits comparable toxicity to the *Anopheles* mosquito [52] but is considered less toxic to fish [53]. We have chosen this compound to assess the impact of target site insensitivity conferring resistance to pyrethroids on SCBIs action on VGSCs. Type I pyrethroids are known to modify resting or inactivated channels, while type II pyrethroids preferably bind to the open state of sodium channels [10]. As etofenprox, a non-ester pyrethroid, represents a separate type, we docked this insecticide to both open and inactivated models of mosquito VGSC.

**Figure 4.**
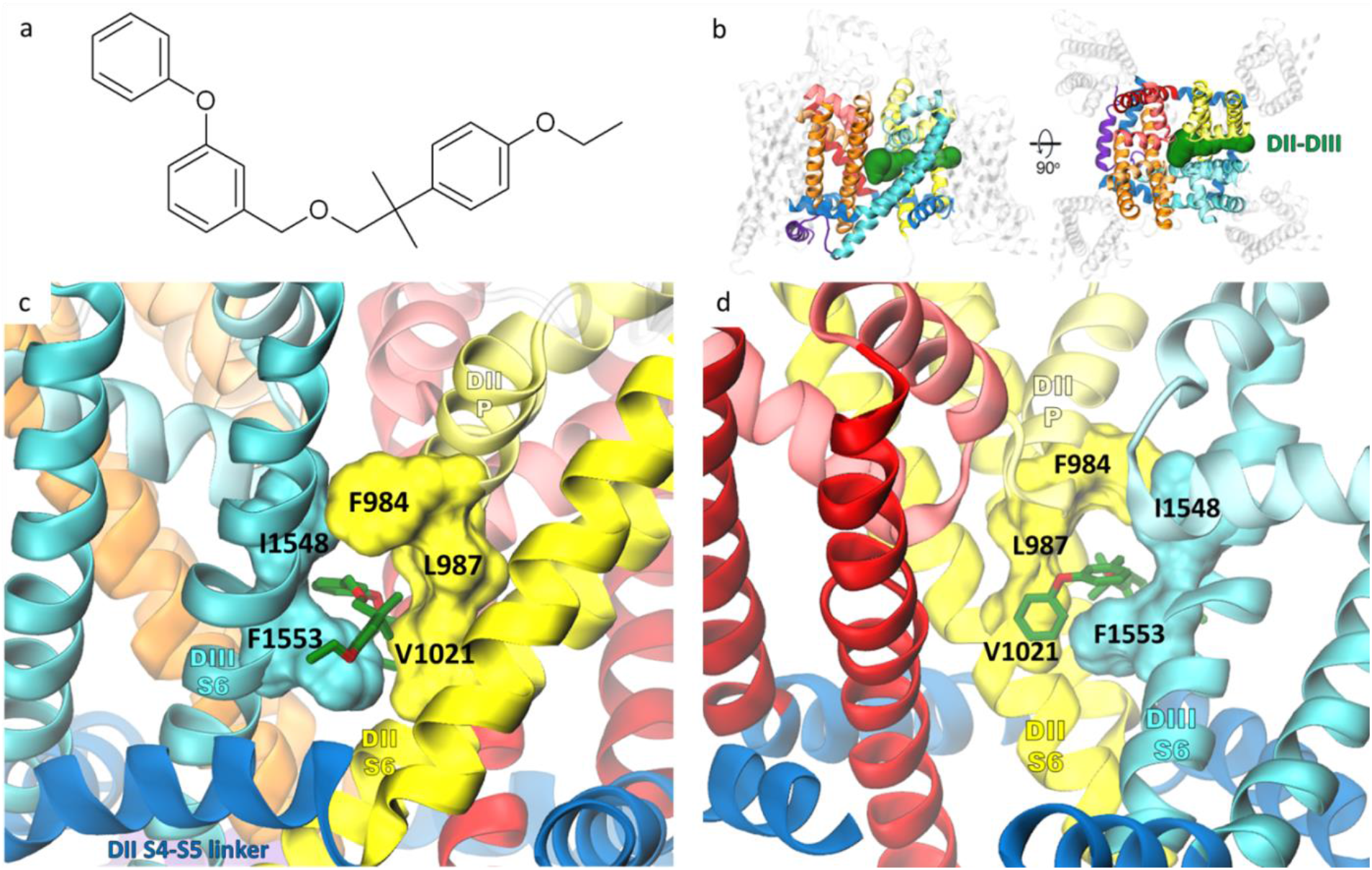
a) Chemical structure of non-ester pyrethroid insecticide etofenprox. b) The DII-DIII fenestration tunnel, found in docking as a ligand binding region, is presented in green in a voltage-gated sodium channel model in a side (left) and a top (right) view. c-d) The lowest energy pose of etofenprox (in green stick representation) bound in the open model. View from the channel/membrane interface is shown in (c), while the fenestration entrance to the central cavity of the channel is visualized in (d), with the PDIII helices removed for clarity. Residues that were found as kdr contributing to both etofenprox and SCBIs binding are shown in a surface representation. The coloring of the channel refers to Figure 1.

The lowest energy poses in the open and inactivated models overlap but are not identical with the SSF values −9.97 kcal/mol and −9.71 kcal/mol, respectively. Etofenprox penetrates the DII-DIII fenestration deeper than it was proposed before in docking to the prokaryotic NavMs-based model [51, 54]. Deep binding results in interactions with 16 kdr residues (Table 1), accounting for 68% and 60% of the total SSF found for the open and inactivated model, respectively. The highest contribution to the binding energy (−2.0 kcal/mol for the open and −2.42 kcal/mol for inactivated model) comes from F1553, which was found as a key bottlenecking residue in the DII-DIII fenestration [25]. In the inactivated-state model, ligand position is stabilized by parallel π-stacking interactions with F1553 and F1025 (that are both kdr, see Table 1). As both overall binding energy and the contribution from the kdr residues were more negative in docking to the open model, we further refer to this pose, presented in Figure 4c,d.

**Table 1.** Etofenprox binding region. DCJW and metaflumizone interactions with given residues are marked with D and M, respectively.

| VGSC domain | *Anopheles gambiae* | *Musca domestica* | Kdr ref. | Overlap with SCBIs binding region |
| --- | --- | --- | --- | --- |
| DII, S4-S5 linker | M923 | M918 | [55-57] | - |
|  | T926 | T921 | - | - |
|  | M927 | M922 | - | - |
| DII S5 | T934 | T929 | [55, 58] | - |
|  | L937 | L932 | [59-62] | M |
|  | C938 | C933 | [55] | - |
|  | I941 | I936 | [61, 63] | - |
| DII P | **F984** | **F979** | **[57, 64]** | **D, M** |
|  | **L987** | **L982** | **[65, 66]** | **D, M** |
|  | C988 | C983 | - | D, M |
| DII S6 | N1018 | N1013 | [67] | M |
|  | **V1021** | **V1016** | **[65, 68-75]** | **D, M** |
|  | L1022 | L1017 | [76] | M |
|  | L1024 | L1019 | - | - |
|  | F1025 | F1020 | [77, 78] | - |
| DIII P | A1510 | A1494 | - | D, M |
|  | F1512 | F1496 | - | D, M |
| DIII S6 | **I1548** | **I1532** | **[75, 79-81]** | **D, M** |
|  | I1549 | I1533 | [82] | D, M |
|  | F1550 | F1534 | [74, 75, 83] | - |
|  | S1552 | S1536 | - | D, M |
|  | **F1553** | **F1537** | **[82, 84]** | **D, M** |
|  | F1554 | F1538 | [85, 86] | - |
|  | L1556 | L1540 | - | D, M |

We found 10 residues to contribute to etofenprox and both SCBIs binding (Table 1). Additional four are shared in etofenprox and metaflumizone but not in the DCJW docking region. Mutation of five residues participating in the binding of all three insecticides (F984, L987, V1021, I1548, and F1553) are found in pyrethroid-resistant insects. We show them in a surface representation in Figure 4c,d. The contribution to the binding energy from kdr residues is higher for metaflumizone than indoxacarb (24% and 15% of total SSF, respectively).

V1021I/G (V1016 in house fly VGSC) is one of the most frequently reported kdr mutations in pyrethroids-resistant insects worldwide. It has been found in disease-spreading *Aedes* mosquitoes in Indonesia [65], Thailand [68, 69], Taiwan [70], Singapore [71], China [72], and Latin America [73]. In mosquitoes, a concomitant S989P+V1016G kdr mutation confers resistance to 17 structurally diverse pyrethroids with the highest level (107 fold) to etofenprox. Cross-resistance to both indoxacarb and DCJW was reported in this study [23]. In *Aedes aegypti* mosquito, the level of resistance to DCJW conferred by the *kdr* allele 410L+1016I+1534C (house fly numbering) was higher than in mosquitoes carrying the 1534C allele alone [87], suggesting the direct role of the V1016I mutation on DCJW action on VGSC. The lysine substitution of the corresponding residue in the rat rNav1.4 channel (V787K) completely abolished mutated VGSC inhibition by metaflumizone [39]. To test the impact of V1021 substitution on SCBI binding, we performed docking to the following VGSC mutants: V1021I, V1021G, and V1021K.

Substitution of V1021 did not affect DCJW binding significantly – RMSD values for a ligand shifted by 0.16 Å and 1.0 Å for isoleucine and glycine, respectively, which are kdr mutations found in nature. Only the lysine substitution pulled the trifluoromethyl group of DCJW out of the CC entrance to the DII-DIII fenestration (RMSD=2.28 Å), reducing the binding affinity by 0.55 kcal/mol (Table 2). A higher effect was observed for metaflumizone. While isoleucine and glycine substitutions only slightly shifted the ligand positions (RMSD ~1.7 Å), in the V1021K substituted channel WT-like binding poses were not found. Our results are in agreement with the previous mutagenesis study on the rNav1.4 channel, where sensitivity to inhibition by 10 μM metaflumizone was abolished entirely in the rNav1.4/V787K channels, but no significant effect was observed for DCJW [39]. V1021K substitution disabled etofenprox binding deep into the DII-DIII fenestration. In this mutant, we obtained a binding pose similar to that presented before [54], with the SSF reduced by 1.7 kcal/mol when compared to the WT (SI Figure S4).

**Table 2.** The affinity of insecticides to the V1021 substituted VGSC. Binding energy calculated with smina scoring function is shown in kcal/mol.

| VGSC | DCJW | metaflumizone | etofenprox |
| --- | --- | --- | --- |
| WT | -9.61 | -10.16 | -9.97 |
| V1021I | ≈ | -9.96 | -9.30 |
| V1021G | -9.38 | -9.66 | -8.99 |
| V1021K | -9.06 | no WT-like poses | -8.30 |

V1021 clearly contributes to SCBI binding. However, neither the isoleucine nor glycine substitution found in resistant insects affects ligand binding to the extent that could substantially decrease the inhibitory effect on mutated channels. As lysine substitution itself enhances slow inactivation [39], the V1021K mutant should not be expected in insects.

## 3. CONCLUSIONS

In this work, we found the binding poses of DCJW and metaflumizone located in the CC of mosquito VGSC, where they interact with residues known to affect channel sensitivity to SCBIs. We analyzed the possible pathways of DCJW entrance to its binding site and found the DIII-DIV fenestration to be the most probable route. The phenylalanine at the midpoint of the DIV S6 helix (F1852), mutated to tyrosine in SCBI-resistant insects, bottlenecks the fenestration and play a key role in stabilization of the ligand in its binding site. Based on our docking and metadynamics studies we postulate that the reduced sensitivity to DCJW in resistant insects, and the increased toxicity of DCJW when compared to its bulkier parent compound, indoxacarb, can be explained by the impeded access of the ligands to their binding site in the CC. Furthermore, by docking etofenprox, we delineated residues that contribute to SCBI and pyrethroid binding sites simultaneously and thus could be responsible for target site cross-resistance. Our study is a step towards better understanding the action of SCBIs on insect VGSCs that should facilitate the fight against insect vector-borne diseases.

## 4. METHODS

### 4.1 Homology Modeling

The homology models of an α subunit of the *Anopheles gambiae* sodium channel protein AgNav1 in open, inactivated, and closed states were built using the SWISS-MODEL server [88], using rat rNav1.5 (PDB: 7FBS [32]), human hNav1.7 (PDB: 6J8G [11]), and cockroach NavPas (PDB code: 6A90 [33]) crystallographic structures, respectively, as templates. The models were built based on the A5I843_ANOGA amino acid sequence provided by the UniProtKB database [89]. Two long, disordered intracellular loops (ICL1 – 338 residues, ICL2 – 240 residues), not present in any sodium channel protein experimental structures, were removed from each model. The quality of the models was validated using ERRAT [90] and PROCHECK [91]. The protein preparation module of Schrodinger Maestro [92] was used to assign protonation states, add hydrogen atoms, and minimize all three models. 95.7% of the inactivated model residues fall below the 95% rejection limit in the ERRAT analysis. Only 8 residues were found in disallowed regions by PROCHECK, all of them belonging to the intracellular or extracellular loops. Mutations of V1021 were introduced using Schrodinger Maestro, followed by minimization of mutated models.

### 4.2 Docking

3D structures of all ligands were downloaded from PubChem [93] and minimized using Schrodinger Maestro LigPrep. Molecular docking was performed using the smina package [94], a fork of Autodock Vina [95] that provides enhanced support for minimization and scoring. Ten independent runs (yielding up to 100 poses each) of flexible ligand docking to each model with default parameters were conducted.

### 4.3 Equilibrium MD Simulations

Inputs for equilibrium MD simulations were generated using the CHARMM-GUI membrane builder [96–98]. To mimic an insect-like membrane, a heterogeneous bilayer model composed of 500 lipid molecules in proportions: 38% DOPE, 18% DOPS, 16% DOPC, 13% POPI, 11% SM (CER180), 3% DOPG, and 1% PALO 16:1 fatty acid was built as proposed for an insect muscarinic GPCR before [99]. Water molecules in the TIP3P model were added above and below the lipids to generate a 20 Å thickness layer further neutralized with Na^+^ and Cl^-^ ions to the concentration of 0.15 M. Six steps of equilibration simulations in the NVT ensemble followed by the NPT ensemble with gradually decreased restraint force constants to various components were run using NAMD 2.13 [100] with the CHARMM36 force field applied. Three independent simulations of 250 ns each were run for the inactivated-state model. Temperature, controlled by the Langevin thermostat, was set to 303.15 K and pressure to 1.01325 bar (1 atm). All MD simulations employed a 2 fs time step.

The MD trajectories were processed into protein-only PDB snapshots saving a frame every 1 ns. DCJW and metaflumizone were docked to each snapshot as described above for the static models, generating up to 750000 poses in total. The lowest energy poses were selected for the starting points of MD simulations of ligand-bound protein. Equilibration followed by 250 ns MD simulations were run as described for the apo protein. Topology and parameters files for DCJW were generated by SwissParam [101]. Results were visualized with the VMD code [102].

### 4.4 Enhanced Sampling MD Simulations

Enhanced sampling MD simulations were run using the Gromacs 2020.7 software [103] patched with the PLUMED 2.8 plugin [104, 105]. The simulations were run in the NVT ensemble using the stochastic velocity rescaling thermostat [106] at 303.15 K with a relaxation time of 1 ps. Hydrogen bonds were constrained using P-LINCS [107]. The voltage-sensor domains of VGSC were removed and neutral groups were added to cap S4-S5 linkers termini. The system preparation (e.g., equilibration) and the rest of simulation parameters were the same as for equilibrium MD simulations (Section 2.3).

To find possible reaction coordinates for DCJW binding in the fenestrations, we used the maze module of PLUMED [108] and followed the protocol described in Refs. [41, 108]. DCJW was biased to move with a constant velocity of 10 Å/ns with a bias height of 240 kcal/mol. The direction of biasing was found by minimizing a loss function describing contacts between the ligand and VGSC every 1 ns using simulated annealing. The implemented loss function was *Q*= Σ_kl_[1+exp(r_kl_-r_0_)]^-1^, where r_kl_ is the distance between the i-th atom of DCJW and the j-th atom of VGSC and r_0_ is set to 4 Å. In total, 30 such MD simulations were run. The simulations were terminated when DCJW dissociated from VGSC or the simulation time exceeded 100 ns. The average length of these MD simulations was around 23 ns.

To reconstruct free-energy profiles along the identified reaction coordinates, for each, we ran a 500-ns well-tempered metadynamics simulation [109]. DCJW conformations and VGSC Cα atoms within 8 Å of any atom of DCJW taken from the reaction coordinates were used to define path-collective variables—progress along the reaction coordinate (s) and the distance from the reaction coordinate (z) [42]. To prohibit DCJW from escaping to the membrane, additional constraints for the progress along the reaction coordinate and the distance from the reaction coordinate with force constants equal to 1000 kJ/mol were added. Both path-collective variables were biased using an initial Gaussian height of 2 kJ/mol, a width of 0.12, and a bias factor of 50. Gaussians were deposited every 1 ps, and the time-dependent constant c(t) was updated every 10 Gaussian depositions. The free-energy profiles were reweighted using the Tiwary-Parrinello algorithm [110], taking into account the metadynamics bias potential and the constraints.

### 4.5 Fenestration Analysis

Fenestration analysis was performed using CAVER 3.0 [43] with default parameters, except for a probe radius of 0.8 Å, shell radius of 15 Å, and shell depth of 15 Å. Three MD trajectories of the inactivated model were processed into a series of PDB snapshots containing pore domain and S4-S5 linkers only, taking a frame every 1 ns. The following residues were used as starting points for tunnel search: Phe416 (S6 DI), Gly1015 (S6 DII), Phe1553 (S6 DIII), and Phe1852 (S6 DIV).

## Supporting Information Available

Supporting Information is available free of charge at XXXXXX.

- Docking of DCJW and metaflumizone to the closed-state AgNav1
- DCJW interaction with sodium ion
- DCJW-channel contacts found in MD simulations
- Docking of indoxacarb
- Docking of etofenprox to the V1016K mutant

## Acknowledgements

This research was funded by the National Science Centre, Poland, under grants no. 2021/41/N/NZ3/02165 (BN) and Sonata 2021/43/D/ST4/00920 (JR). J.R. acknowledges funding from the Polish Science Foundation (START) and the Ministry of Science and Higher Education in Poland. Simulations were carried out using computing clusters at the Centre of Informatics Tricity Academic Supercomputer & Network and the Interdisciplinary Centre for Mathematical and Computational Modeling (ICM), University of Warsaw, under grant no. GA76-16. Facilities of UMK/ICNT were used during this project. BN, JR, and WN are members of #MEMO-BIT and EF IDUB UMK teams.

## Contributions

BN: Conceptualization; Data curation; Formal analysis; Investigation; Methodology; Resources; Software; Validation; Visualization; Writing - original draft JR: Data curation; Formal analysis; Investigation; Methodology; Software; Validation; Writing - review & editing BL: Conceptualization; Supervision; Writing - review & editing WN: Conceptualization; Funding acquisition; Project administration; Resources; Supervision; Validation; Writing - review & editing

