## Supporting Information for "Toward overcoming pyrethroid resistance in mosquito control: the role of sodium channel blocker insecticides"

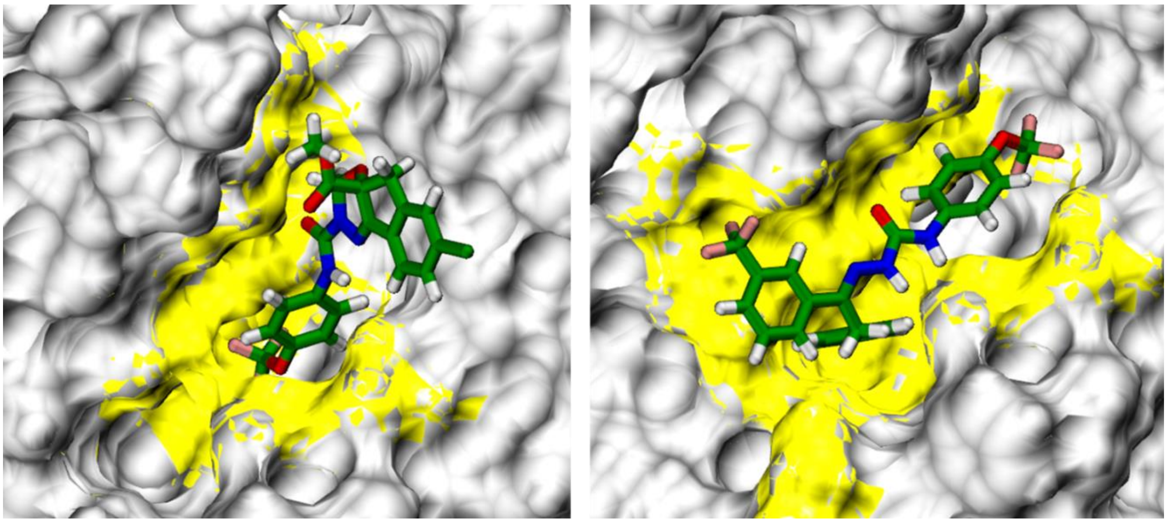

DCJW, SSF=-8.05 kcal/mol

metaflumizone, SSF=-8.42 kcal/mol

Figure S1. Docking of DCJW (left) and metaflumizone (right) to the closed-state AgNav1 model. In their lowest energy poses (SSF, smina scoring function) ligands approach the DIII-DIV fenestration. Hydrophobic residues in a distance of 5 Å from each ligand are in yellow.

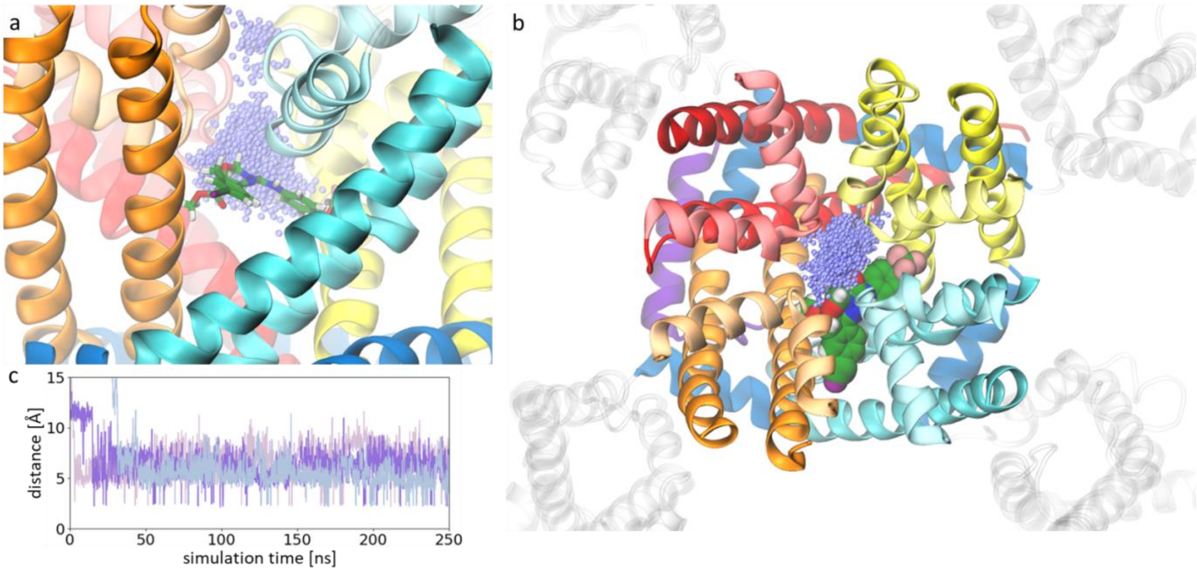


Figure S2. DCJW interaction with sodium ion. Side (a) and top (b) view of the inactivated-state mosquito VGSC with DCJW bound. The sodium ions positions found in the pore domain of the channel in three 250 ns MD trajectories are overlapped and presented as violet dots. The shorted distance between the ion and the carbonyl group of DCJW is shown in (c) where three shades of violet indicate three MD trajectories.

SI Table 1. Percentage of MD snapshots in which contact between DCJW and a given VGSC residue was found. Residues known to affect channel sensitivity to SCBIs are bolded.

| segment | residue | MD1 | MD2 | MD3 | mean |
|---|---|---|---|---|---|
| S6 DIV | **F1852** | 94 | 97 | 97 | 96 |
| S6 DIII | F1553 | 81 | 91 | 91 | 88 |
| DIII P | F1512 | 73 | 94 | 83 | 83 |
| DIII P | T1511 | 77 | 76 | 87 | 80 |
| S6 DIII | I1548 | 80 | 83 | 75 | 79 |
| S6 DIII | L1556 | 78 | 75 | 82 | 78 |
| DII P | C988 | 76 | 75 | 73 | 75 |
| S6 DII | **V1021** | 69 | 75 | 76 | 73 |
| DIII P | A1510 | 73 | 49 | 97 | 73 |
| S6 DIII | S1552 | 64 | 36 | 79 | 60 |
| DIII P | **F1507** | 58 | 73 | 22 | 51 |
| S6 DIII | I1549 | 47 | 50 | 51 | 50 |
| DII P | L987 | 47 | 33 | 58 | 46 |
| S6 DIV | L1848 | 27 | 90 | 12 | 43 |
| S6 DIII | **T1555** | 49 | 00 | 73 | 41 |
| S5 DIII | W1445 | 50 | 01 | 69 | 40 |
| S5 DIII | C1441 | 37 | 00 | 81 | 39 |
| S6 DIV | V1849 | 18 | 84 | 06 | 36 |
| DIV P | S1805 | 26 | 72 | 06 | 34 |
| S5 DIII | L1438 | 32 | 00 | 67 | 33 |
| S6 DIV | **V1855** | 10 | 33 | 01 | 15 |
| S6 DIV | **Y1859** | 11 | 25 | 00 | 12 |
| DII P | F984 | 19 | 09 | 02 | 10 |
| DIV P | **T1804** | 04 | 25 | 00 | 10 |

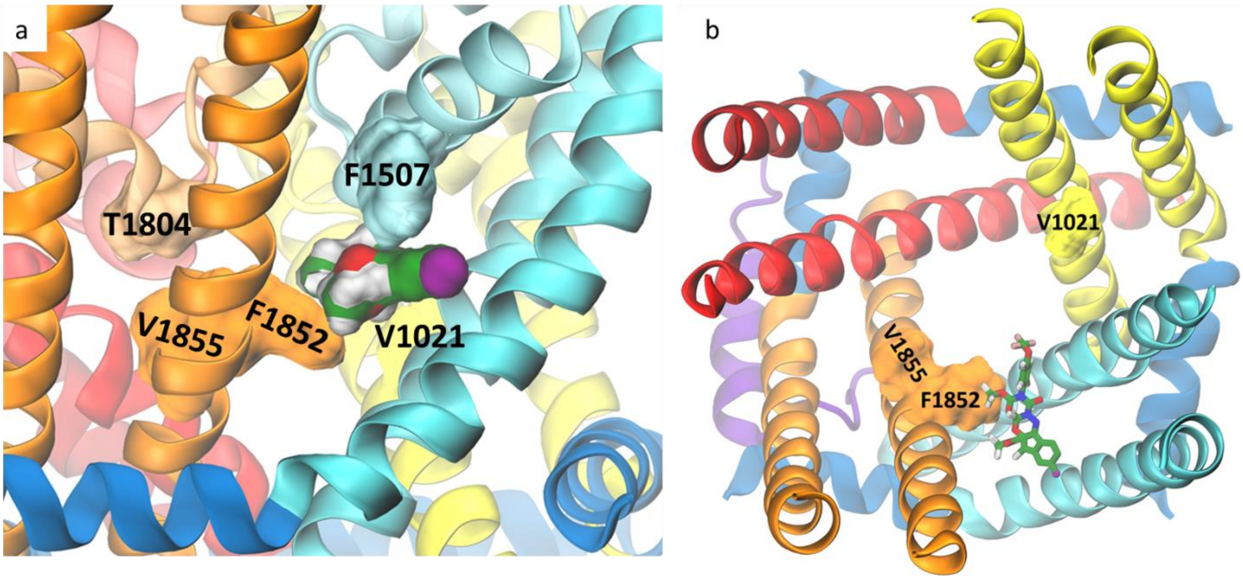


Figure S3. Docking of indoxacarb to the inactivated-state mosquito VGSC. Side view (a) and top view with the P-loops removed for clarity (b) are presented with the residues found to affect the channel sensitivity to blocker insecticides shown in the surface representation.

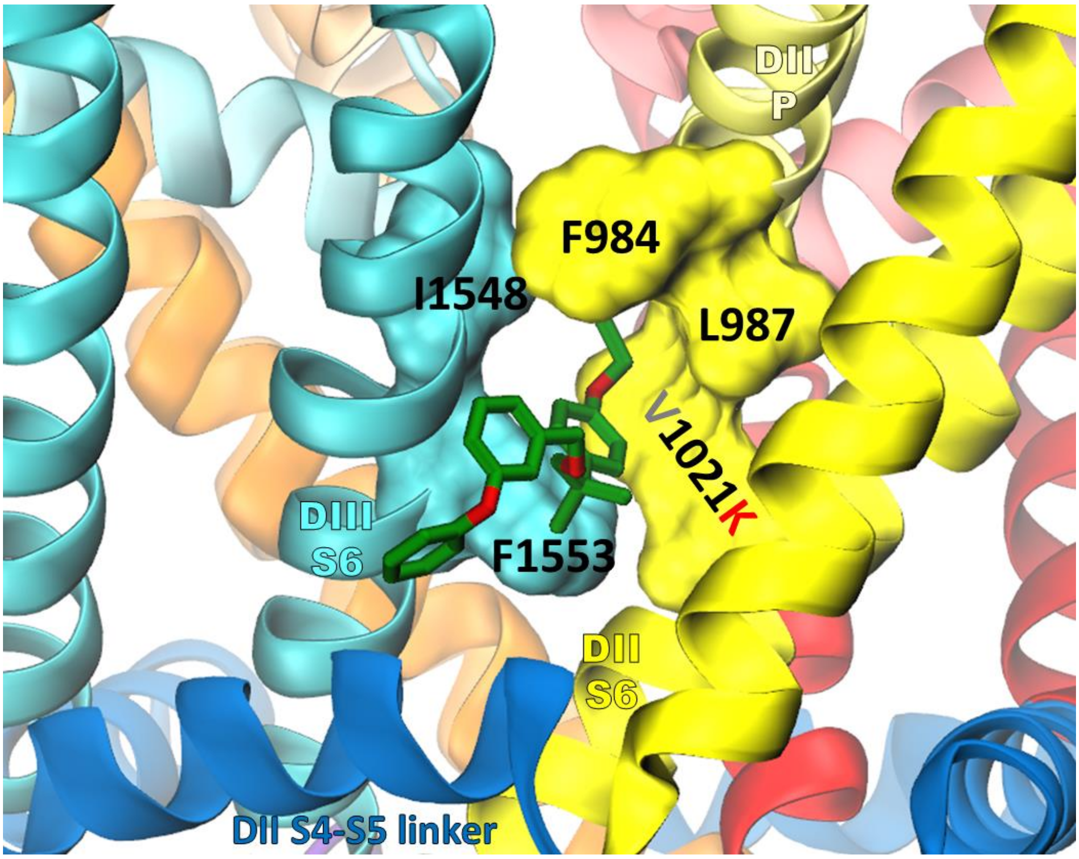


Figure S4. Etofenprox docking to the V1021K mutant of mosquito sodium channel. Residues that were found as kdr contributing to both etofenprox and SCBIs binding are shown in a surface representation.
